# Selective Display of a Chemoattractant Agonist on Cancer Cells Activates the Formyl Peptide Receptor 1 on Immune Cells

**DOI:** 10.1101/2021.09.27.462035

**Authors:** Eden L. Sikorski, Janessa Wehr, Noel J. Ferraro, Marcos M. Pires, Damien Thévenin

**Affiliations:** Department of Chemistry, Lehigh University, Bethlehem, Pennsylvania, USA; Department of Chemistry, University of Virginia, Charlottesville, Virginia, USA

## Abstract

Current immunotherapeutics often work by directing components of the immune system to recognize biomarkers on the surface of cancer cells to generate an immune response. However, variable changes in biomarker distribution and expression can result in uneven patient response. The development of a more universal tumor-homing strategy has the potential to improve selectivity and extend therapy to cancers with decreased expression or absence of specific biomarkers. Here, we designed a bifunctional agent that exploits the inherent acidic microenvironment of most solid tumors to selectively graft the surface of cancer cells with a formyl peptide receptor ligand (FPRL). Our approach is based on the pH(Low) Insertion Peptide (pHLIP), a unique peptide that selectively targets tumors *in vivo* by anchoring onto cancer cells in a pH-dependent manner. We establish that selectively remodeling cancer cells with a pHLIP-based FPRL activates formyl peptide receptors on recruited immune cells, potentially initiating an immune response towards tumors.

## Introduction

The first line of defense against infections is the detection of invading pathogens by pattern recognition receptors (PRRs) on the surface of innate immune cells. The family of PRRs have evolved to identify infectious agents by their distinct structural and biochemical features, called pathogen-associated molecular patterns (PAMPs).^[1-4]^ One of these unique structural fingerprints is *N*-formyl methionine, which is found in proteins of bacterial pathogens and mitochondrial proteins of ruptured host cells. ^5,6]^ This minor, but structurally discernable, difference enables the immune system to specifically detect the presence of bacteria^[7]^ and respond to infection-associated inflammation^[8]^ via pattern recognition receptors, including a family of G protein-coupled receptors primarily expressed in phagocytic leukocytes (neutrophils, monocytes, dendritic cells, and natural killer cells) known as N-Formyl Peptide Receptors (FPR). Stimulation of FPRs triggers classical GPCR signaling cascades that result in chemotaxis, phagocytosis, cytokine production, and the production of reactive oxygen species.^[6,9]^ While FPRs were originally named after their ability to detect *N*-formylated peptides, a variety of diverse agonists have since been identified.^[10-13]^ For instance, in addition to *N*-formylated peptides, *C*-amidated and unmodified peptides from different origins, as well as several non-peptide ligands can activate FPRs.^[10,12,14]^

Because they are potent chemo-attractants for immune cells,^[15]^ FPR ligands (FPRLs) have tremendous therapeutic potential as immune stimulatory molecules. Consequently, FPRLs have started to be administered to initiate antitumor immune responses.^[16-18]^ For instance, the use of bifunctional synthetic agents composed of a FPRL and a tumor-homing moiety targeting an over-expressed membrane protein (e.g., integrin and specific cancer antigens) has successfully increased immune cell infiltration into tissue and led to tumor shrinkage *in vivo*.^[19-21^

However, prior targeting approaches are limited by their reliance on surface proteins with variable changes in expression level as well as a lack of universal targetable tumor biomarkers among all patients. In contrast, we recently reported a unique class of immunotherapeutic agents that selectively graft epitopes onto the surface of cancer cells based on a potentially universal feature of solid tumors: their inherent acidic microenvironment.^[22]^ Indeed, the microenvironment surrounding nearly all tumor masses regardless of their tissue or cellular origin is acidic (pH 6.0-6.8) in contrast to healthy tissues (pH 7.2-7.5).^[23-28]^ Therefore, the acidic microenvironment of tumors provides a window for selectively targeting tumor masses while sparing healthy tissues. Our tumor-tagging strategy is based on the pH(Low) Insertion Peptide (pHLIP), a peptide with established tumor targeting and exciting therapeutic potential that anchors onto the surface of cancer cells in a pH-dependent manner.^[29-37]^

Remarkably, pHLIP exists as a soluble monomer in neutral aqueous solutions but, under mild acidic conditions (such as in the tumor microenvironment), inserts unidirectionally into the membrane insertion (i.e., extracellular N-terminus) as a transmembrane *α*-helix. This insertion topology can therefore be used to graft a variety of molecules onto cancer cell surfaces as long as they are conjugated to the N-terminus of pHLIP.^[22,38,39]^

Here, we show that remodeling the surface of cancer cells with a FPR ligand using a pHLIP-based targeting strategy could selectively engage an immune response towards tumors by activating FPR1 on recruited immune cells (Figure 1).

**Figure 1.**
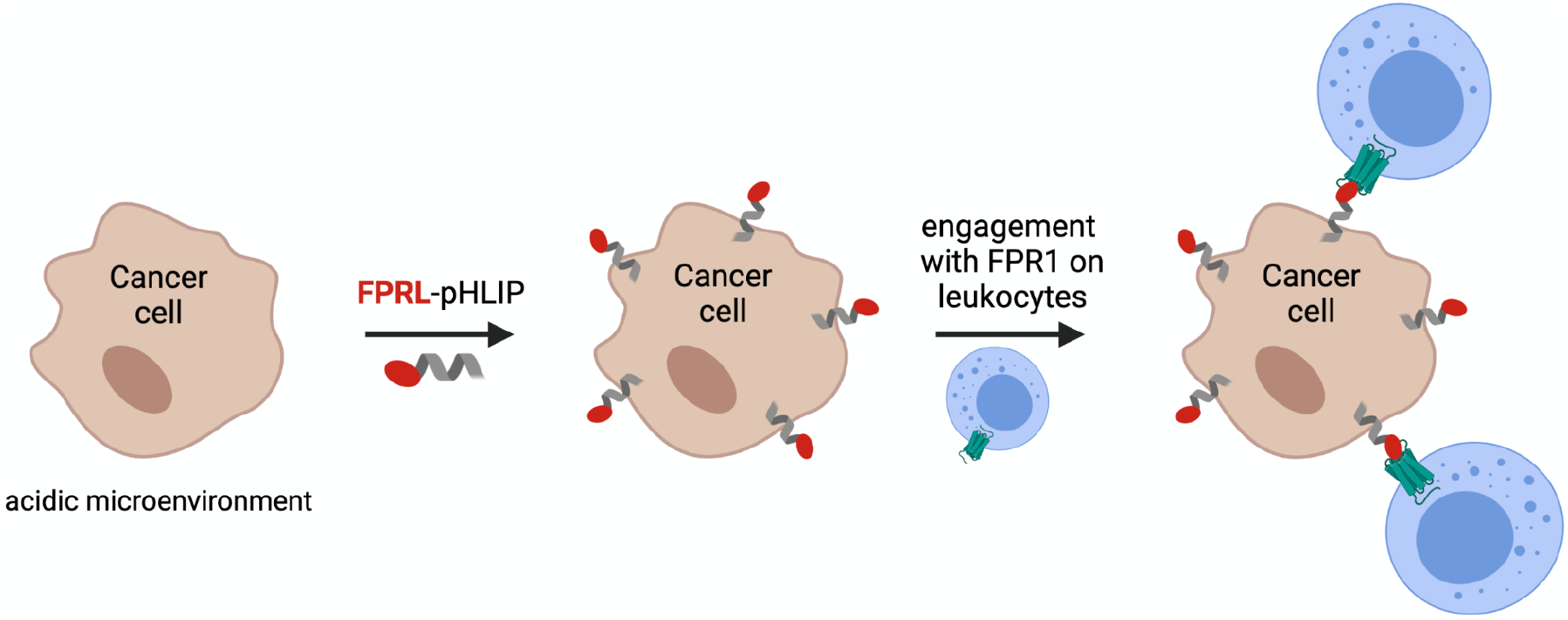
Schematic representation of the mechanism of action of the FPRL-pHLIP conjugate. (created with BioRender.com)

## Results and Discussion

### Design and Synthesis of FPRL-pHLIP Conjugates

While many FPR1 agonists have been identified, we chose to incorporate in our design an hexapeptide with a C-terminal amidated D-methionine (WKYMVm; named in this study, W_D_)^[40]^ because of (i) its small size, (ii) reported structure-function studies, and (iii) demonstrated potency.^[41-43]^ We also used, as a control, the corresponding stereoisomer peptide in which the C-terminal D-methionine is replaced with a *L*-methionine (named in this study, W_L_), -a substitution that has been shown to greatly decreases the peptide biological activity.^[44]^

We conjugated the W-peptides (Table 1), modified with a maleimide-based linker, to the N-terminus of pHLIP the sulfhydryl of a cysteine residue to produce W_D_-pHLIP and W_L_-pHLIP (Scheme 1). Lysine within WKYMVm was chosen as the site of conjugation to pHLIP based on prior derivatives, in which fluorescein isothiocyanate can be conjugated to the lysine without significantly disrupting activation potential of the peptide.^[45]^ Moreover, this linking strategy should display the W peptides to the extracellular face of the plasma membrane of cancer cells upon insertion of pHLIP in an acidic environment, such as the one found in tumours, allowing for the W_D_ peptide to interact with and activate FPR1 present at the surface of immune cells. The FPRL-pHLIP conjugates were purified by RP-HPLC and their identity was confirmed by MALDI-TOF mass spectrometry (Figure S1).

**Table 1.** Sequences of peptides used in this study.

| Conjugate | Sequence |
| --- | --- |
| W<sub>L</sub> | WK(mal)YMVM |
| W<sub>D</sub> | WK(mal)YMV<sub>m</sub> |
| pHLIP | GCEQNPIYWARYADWLFTTPLLLLDLALLVDADEGTG |
| W<sub>L</sub>-pHLIP | GC(WKYMVM)EQNPIYWARYADWLFTTPLLLLDLALLVDADEGTG |
| W<sub>D</sub>-pHLIP | GC(WKYMV<sub>m</sub>)EQNPIYWARYADWLFTTPLLLLDLALLVDADEGTG |

### Interaction of FRPL-pHLIP Conjugates with Lipid Bilayers

Tryptophan fluorescence emission (pHLIP contains two tryptophan residues) and far-UV circular dichroism (CD) spectroscopy were used to determine whether conjugation of W_L_ or W_D_ to pHLIP alters its pH-mediated insertion into lipid bilayers. CD was used to monitor structural changes in the presence of large unilamellar POPC lipid vesicles at both physiological and acidic pH (Figure 2A,C). Both constructs, W_L_-pHLIP and W_D_-pHLIP, exhibit greater *α*-helicity when the pH is decreased from pH 7.4 (blue line) to pH 5.0 (red line), which is characteristic of pHLIP’s pH response. The insertion of the W-pHLIP conjugates into the vesicles was monitored using fluorescence spectroscopy, as the emission of tryptophan is sensitive to the polarity of the local environment (Figure 2B,D). Lowering the pH from 7.4 (blue line) to pH 5.0 (red line) results in a characteristic blue shift and an increase in fluorescence emission intensity for both W_D_-pHLIP and W_L_-pHLIP. These observations reflect the changing environment of the tryptophan residues of pHLIP, switching from being exposed to a polar, aqueous environment to becoming buried in a more hydrophobic area upon pHLIP insertion in the lipid bilayer. These findings are also consistent with the conformational change observed by CD. Taken together, fluorescence and CD measurements demonstrate that the presence of W_L_ or W_D_ at the N-terminus of pHLIP does not significantly disrupt the pH-mediated structural behavior of pHLIP.

**Figure 2.**
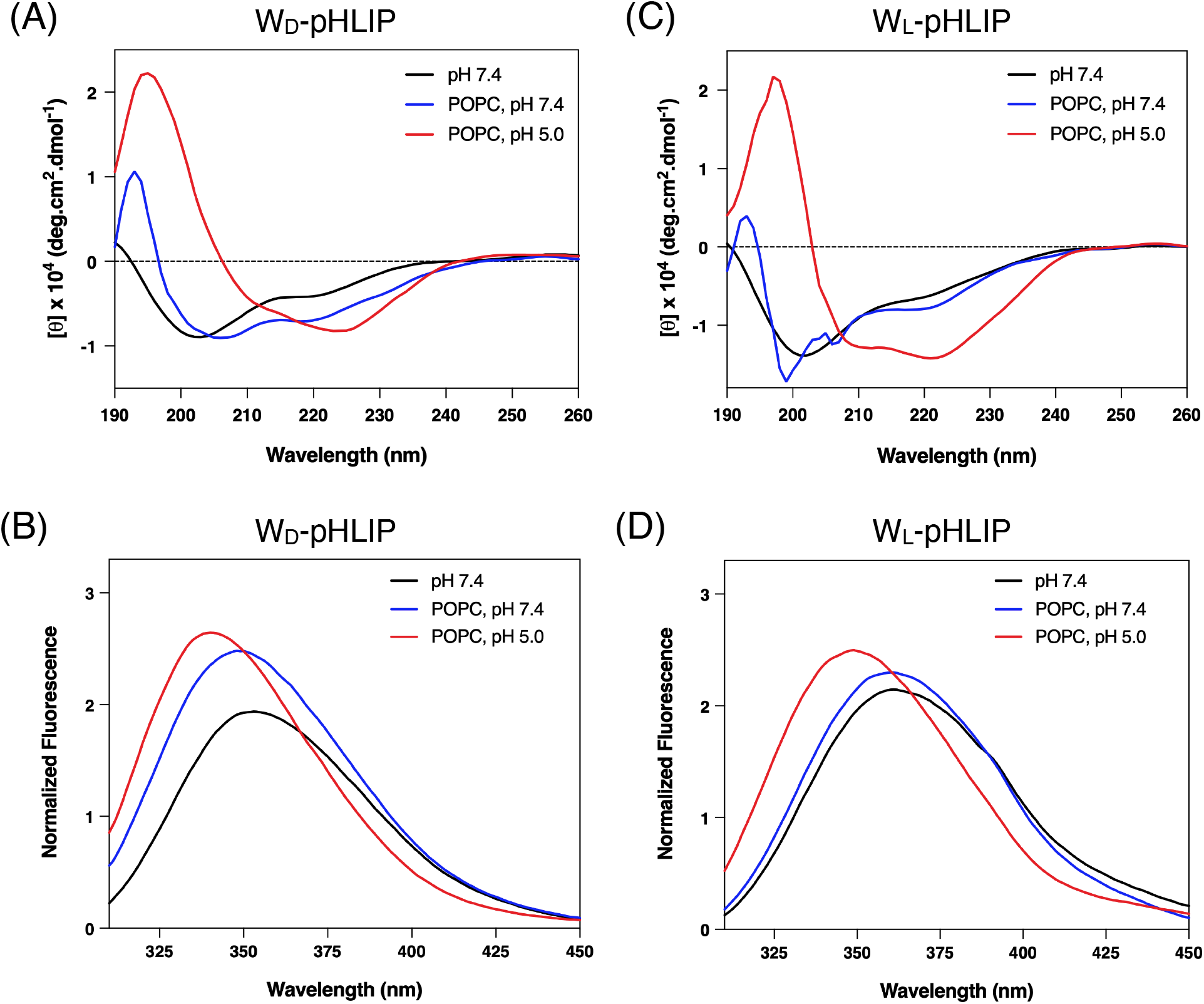
Interaction of W_D_-pHLIP and W_L_-pHLIP with lipid bilayers. Circular dichroism (A,B) and tryptophan fluorescence emission (C,D) in the absence (black) or presence of large unilamellar POPC lipid vesicles at pH 7.4 (blue) and pH 5.0 (red).

### W_D_-pHLIP Activates FPR1 in a pH-Dependent Manner

Prior to testing the ability of the FPRL-pHLIP conjugates to selectively activate FPR1, we confirmed that modifying the W-peptides with a maleimide did not change their biological activity. To characterize the agonistic abilities of the modified W-peptides, the PRESTO-Tango ß-arrestin recruitment assay was used as previously described.^[46,47]^ Briefly, the gene coding for FPR1 fused to the transcriptional activator, tTA, was transfected into HLTA cells stably expressing a tTA-dependent luciferase reporter gene and ß-arrestin-TEV fusion gene. Upon ligand-dependent activation, FPR1 undergoes a conformational change and becomes phosphorylated at specific serine and threonine residues at the cytoplasmic C-terminus resulting in recruitment of ß-arrestin-TEV. The TEV protease cleaves tTA from FPR1 allowing tTA to enter the nucleus and activate expression of the luciferase reporter gene.

The transfected HTLA cells were treated with increasing concentrations of W_D_ or W_L_ for at least 18 hours to allow for receptor activation and signal amplification. Following incubation, the cells were lysed and luminescence was quantified. A concentration-dependent increase in luminescence was observed for both isomers (Figure S2), suggesting that the maleimide modification does not disrupt the ability to activate FPR1. As expected, the stereoisomer control (W_L_) displayed lower activation ability of FPR1 as higher concentrations of agonist were required to elicit a response (Figure S2, dashed line). These results are consistent with previous studies indicating that FPR1 has a stereoselective preference for the D-versus the L-isoform.^[43]^

Next, we evaluated the ability of the W-pHLIP conjugates to selectively insert and decorate the surface of triple-negative MDA-MB-231 breast cancer cells and activate FPR1. Briefly, MDA-MB-231 cells were treated with W_D_-pHLIP, W_L_-pHLIP, or unconjugated pHLIP at pH 7.4 and pH 6.0. The MDA-MB-231 cells were then incubated with transfected HTLA cells and FPR1 activation was monitored using the PRESTO-Tango ß-arrestin recruitment assay described above. A pH-dependent increase in luminescence was observed following treatment with W_D_-pHLIP (Figure 3), suggesting that W_D_ is selectively displayed on the cancer cell surface by pHLIP and is capable of activating FPR1 on adjacent HLTA cells. As expected, no significant increase in luminescence was observed in cells treated with W_L_-pHLIP or unconjugated pHLIP indicating that W_D_ is necessary for FPR1 activation.

**Figure 3.**
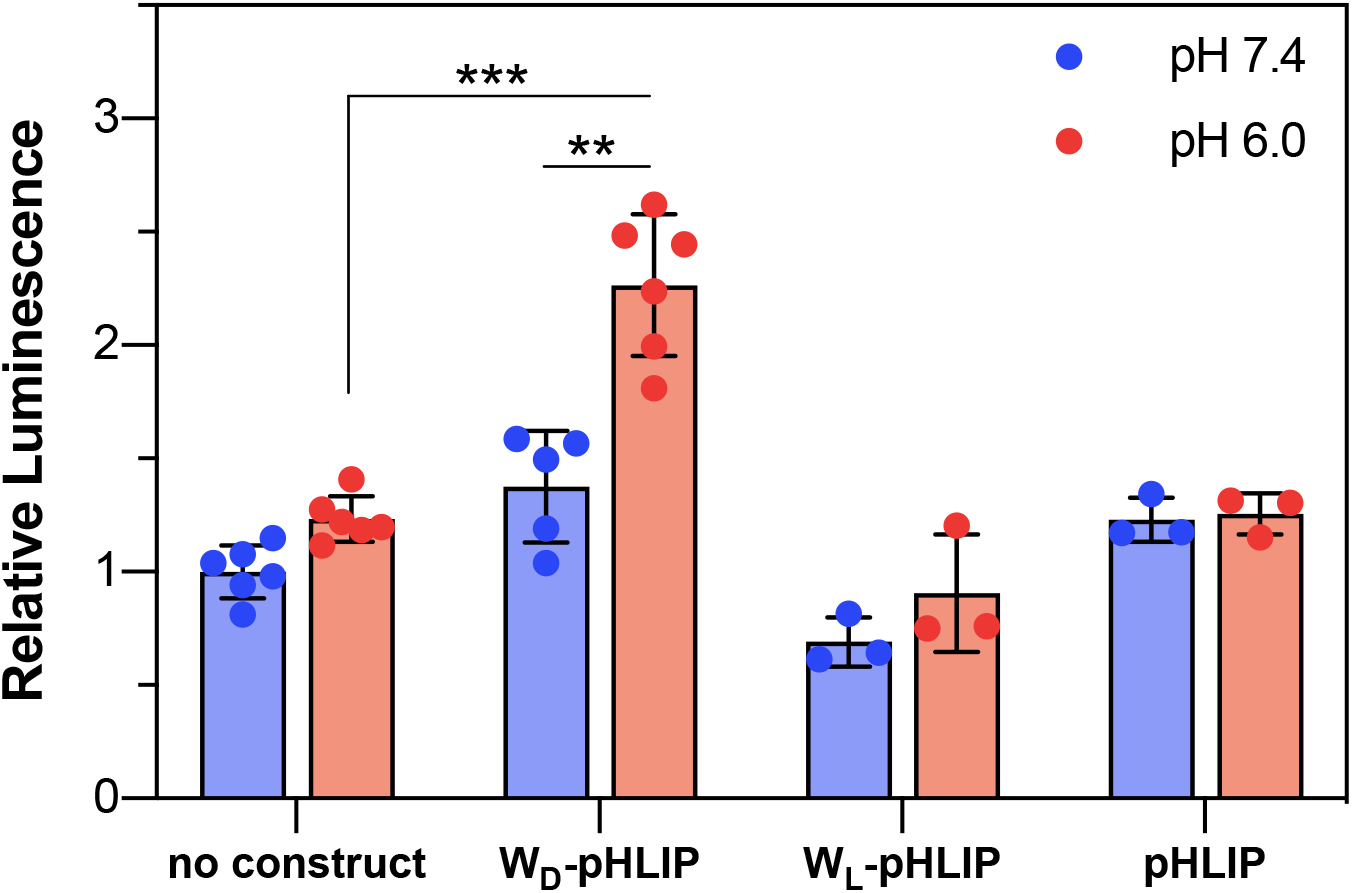
W_D_-pHLIP activates FPR1 in a pH-dependent manner. MDA-MB-231 cells were treated with W_D_-pHLIP, W_L_-pHLIP, or pHLIP at pH 7.4 or 6.0 and incubated with HTLA cells transiently expressing FPR1. Receptor activity was quantified by luciferase expression. Relative luminescence represents the fold increase over that of cells incubated at pH 7.4 only. Results are shown as mean ± standard deviation (*n* = 3-6). Statistical significance was assessed using an unpaired *t* test (at 95% confidence intervals): ***p≤0.001 and \*\**p*≤0.01.

### W_D_-pHLIP Induces Selective Activation of FPR1 at the Surface of Natural Killer Cells

Having established that FPR1 was activated by W_D_-pHLIP, we sought to characterize downstream functional effects in native immune cells by monitoring changes in intracellular calcium fluxes. Upon activation, FPR1 stimulates phospholipase Cß via the dissociated ßƔ G-protein subunits leading to the release of calcium from intracellular stores to elicit cell signaling.^[48]^ We monitored intracellular calcium mobilization in natural killer (NK) cells that naturally express FPR1 by using the calcium sensitive indicator fluo-4 that fluoresces when bound to calcium. Briefly, MDA-MB-231 cells were treated with W_D_-pHLIP or W_L_-pHLIP and incubated with NK cells preloaded with fluo-4. An increase in fluorescence intensity was observed in NK cells incubated with MDA-MB-231 cells pretreated with W_D_-pHLIP at pH 6.0 compared to pH 7.4 (Figure 4A,B), indicating intracellular calcium fluxes were caused by selective FPR1 activation. Conversely, no significant changes in fluorescence intensity was observed when NK cells were incubated with MDA-MB-231 cells pretreated with W_L_-pHLIP at either pH 7.4 or 6.0 suggesting minimal FPR1 activation.

**Figure 4.**
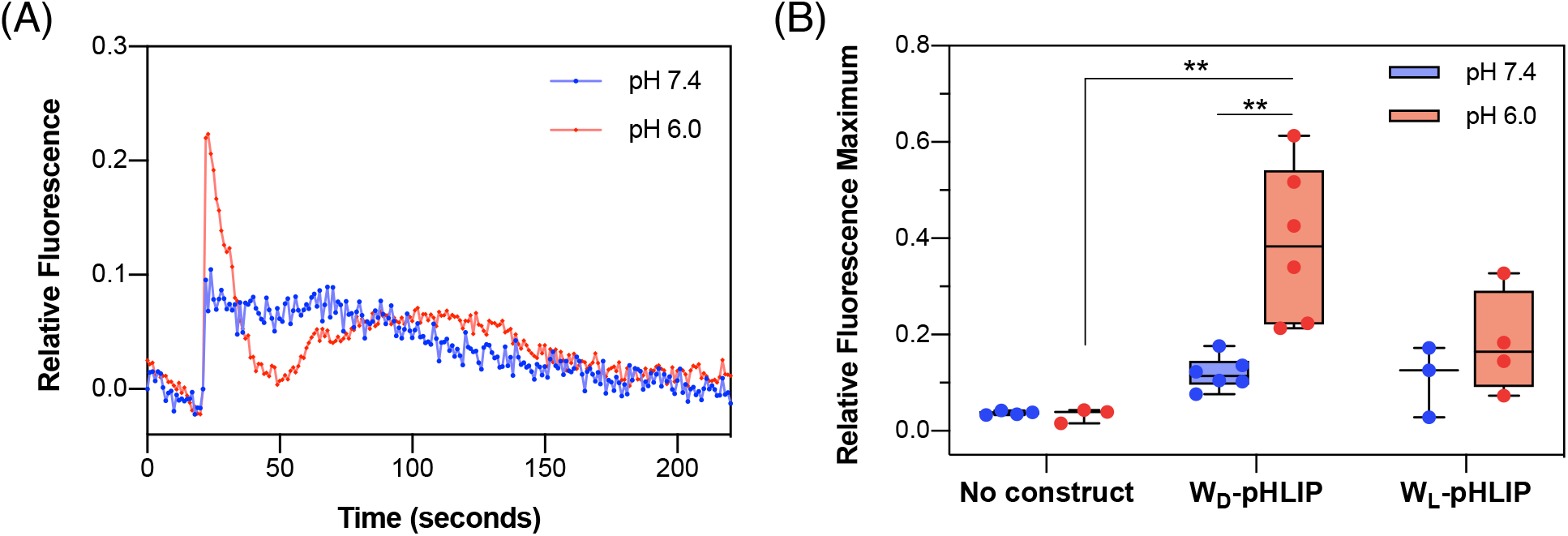
W_D_-pHLIP activates FPR1 on recruited NK cells in a pH-dependent manner. NK cells were loaded with Fluo-4 and incubated with MDA-MB-231 cells treated with FPRL-pHLIP conjugates at pH 7.4 or 6.0. (A) Representative Ca^2+^ fluorescence response profiles upon addition of MDA-MB-231 cells treated with FP_D_-pHLIP at pH 7.4 and 6.0. (B) Quantification of the maximum fluorescence response observed. Relative fluorescence represents the fold increase over baseline. Results are shown as mean ± standard error (*n* = 3-6). Statistical significance was assessed using an unpaired *t* test (at 95% confidence intervals): \*\**p*≤0.01.

## Conclusion

We have designed and tested immune-engaging agents that have superior targeting properties towards cancer cells by exploiting the acidic microenvironment of tumors. We show that grafting the surface of cancer cells with a FPRL initiates an immune response by selective activation of FPR1 on recruited immune cells. Considering that the activation of FPR1 through the release of damage-associated molecular pattern molecules at the tumor site appears to be crucial for chemotherapy-induced antitumor immunity,^[49]^ our approach may provide an effective synergistic effect to conventional chemotherapy. Optimization efforts in construct design and further evaluation both *in vitro* and *in vivo* are currently ongoing in our laboratories.

## Experimental Section

### Peptide Synthesis, Purification, and Conjugation

Both isoforms of the W-peptide were prepared by solid-phase peptide synthesis using rink amide resin (Chem-Impex #12662). pHLIP, containing a cysteine residue at its N-terminus, was prepared with a CEM Liberty Blue microwave peptide synthesizer using rink amide resin (CEM 0.19 mmol/g loading capacity). Briefly, each amino acid, used as a 0.2 M solution in DMF, was coupled using DIC or HBTU as the activator and oxyma or DIEA as the activator base. Removal of the Fmoc protecting group after each coupling step was facilitated using 6% piperazine and 0.1 M HOBt in DMF. The side chain lysine residue of both W-peptides were protected with 4-methyltrityl (Fmoc-Lys(MTT)-OH, Chem-Impex #03720). The MTT protection group was selectively removed with several washes of 0.1% TFA in DCM for 15 minutes at room temperature. The free amine of the lysine side chain was coupled to 3-maleimidopropionic acid (2 equivalents, Chem-Impex #14709) using HBTU and DIEA for 2 hours at room temperature (Scheme S1). The peptides were cleaved from the resin and purified via RP-HPLC (Phenomenex Luna Omega, 5 μm, 250 × 21.20 mm C18; flow rate 5 mL/min; phase A: water 0.1% TFA; phase B: acetonitrile 0.1% TFA; 60 min gradient from 95:5 to 0:100 A/B). Following purification, the W-peptides (3 equivalents) in DMF were conjugated to the N-terminal cysteine residue of pHLIP solubilized in 50 mM HEPES, pH 7.4 for 4 hours at room temperature. The resulting conjugates, W_D_-pHLIP and W_L_-pHLIP, were again purified by RP-HPLC as described above. The purity of each conjugate was determined by analytical RP-HPLC (Phenomenex Luna Omega, 5 μm, 250 × 10 mm C18; flow rate 5 mL/min; phase A: water 0.01% TFA; phase B: acetonitrile 0.01% TFA; 60 min gradient from 95:5 to 0:100 A/B), and their identity was confirmed via MALDI-TOF mass spectrometry (Shimadzu 8020) (Figure S2).

### Sample Preparation of CD and Tryptophan Fluorescence Measurements

All peptide constructs were solubilized to 20 µM in 5 mM sodium phosphate (pH 8.0). Each construct was diluted to a final concentration of 7 μM before analysis. For measurements in vesicles, 1-palmitoyl-2-oleoyl-sn-glycero-3-phosphocholine (POPC) was dried as a thin film and held under vacuum for at least 24 hours. The lipids were then rehydrated in 5 mM sodium phosphate, pH 8.0 for at least 30 minutes with periodic gentle vortexing. The resulting large multilamellar vesicles were freeze-thawed for seven cycles and subsequently extruded through a polycarbonate membrane with 100 nm pores using a Mini-Extruder (Avanti Polar Lipids) to produce large unilamellar vesicles (LUVs). The constructs were incubated with the resulting LUVs at a 1:300 ratio. The pH was adjusted to the desired experimental values with HCl, and the samples were incubated for 30 minutes at room temperature prior to spectroscopic analysis.

### Circular Dichroism (CD) Spectroscopy

Far-UV CD spectra were recorded on a Jasco J-815 CD spectrometer equipped with a Peltier thermal-controlled cuvette holder (Jasco). Measurements were performed in 0.1 mm quartz cuvette. CD intensities are expressed in mean residue molar ellipticity [θ] calculated from the following equation: 

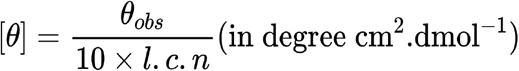

where, θ_obs_ is the observed ellipticity in millidegrees, l is the optical path length in centimeters, c is the final molar concentration of the peptides, and n is the number of amino acid residues. Raw data was acquired from 260 nm to 200 nm at 1 nm intervals with a 100 nm/min scan rate, and at least five scans were averaged for each sample. The spectrum of POPC LUVs was subtracted from all construct samples.

### Tryptophan fluorescence spectroscopy

Fluorescence emission spectra were acquired with a Fluorolog-3 Spectrofluorometer (HORIBA). The excitation wavelength was set at 280 nm and the emission spectrum was measured from 300 to 450 nm. The excitation and emission slits were both set to 5 nm.

### Cell Culture

Human breast adenocarcinoma MDA-MB-231 cells were cultured in Dulbecco’s modified Eagle’s medium (DMEM) high glucose supplemented with 10% fetal bovine serum (FBS), 100 units/mL penicillin, and 0.1 mg/mL streptomycin. Human embryonic kidney cells (HTLA; a HEK293-derived cell line containing stable integrations of a tTA-dependent luciferase reporter and β-arrestin2-TEV fusion gene) were maintained in DMEM supplemented with 10% FBS, 2 μg/mL puromycin, and 100 μg/mL hygromycin. Engineered natural killer NK-92 (haNK cells; ATCC PTA-6967) were cultured in minimum essential medium (MEMα) without nucleosides supplemented with 0.1 mM 2-mercaptoethanol, 0.2 mM inositol, 0.02 mM folic acid, 12.5% FBS, 12.5% horse serum, 100-200 units/mL interleukin-2 (IL-2), 100 units/mL penicillin, and 0.1 mg/mL streptomycin. All cells were cultured in a humidified atmosphere of 5% CO_2_ at 37 °C.

### Tango Assay

HTLA cells stably expressing the tTA-dependent luciferase reporter and β-arrestin-TEV fusion gene (kind gift from Prof. Ryan Roth; University of North Carolina at Chapel Hill) were seeded at a density of 150,000 cells/well in a 6-well plate and incubated at 37 °C to until the cells to reach ∼60% confluency. Transient transfections were performed with FPR1-tTA (Addgene #66283) using calcium phosphate as previously described.{Jordan:1996hf} The following day, the transfected cells were seeded at a density of 40,000 cells/well in 96-well plate pretreated with poly-l-lysine and allowed to adhere overnight at 37 °C. For determining the agonistic activity of the unconjugated W_D_ and W_L_ peptides, the transfected HLTA cells were washed with Hanks Balanced Salt solution (HBSS) and incubated with varying concentrations of W_D_ and W_L_ solubilized in HBSS at 37 °C for at least 18 hours. Luminescence was quantified with an Infinite F200 Pro microplate reader (Tecan) using the Bright-Glo™ Luciferase Assay System (Promega #E2610) according to the manufacturer’s protocol. For determining the agonistic activity of W_D_-pHLIP and W_L_-pHLIP, the conjugates were resuspended in 5 mM sodium phosphate (pH 9.0) to a concentration of 20 µM and incubated at room temperature for 1 hour. Immediately following the incubation, each conjugate was diluted to 12.5 µM by addition of PBS (pH 7.4) so that upon pH adjustment with 10 mM sodium citrate in PBS (pH 2.0), the desired treatment concentration (10 µM) was obtained. MDA-MB-231 cells were harvested and washed twice with PBS (pH 7.4). The MDA-MB-231 cells were then treated in suspension with 10 µM of either conjugate for 10 minutes at 37 °C at pH 7.4 or 6.0. After treatment, the MDA-MB-231 cells were washed once with PBS at the same pH as treatment and resuspended in HBSS. Next, the transfected HTLA cells were washed with HBSS and 5,000 treated MDA-MB-231 cells were added to each well. The cells were incubated together for at least 18 hours at 37 °C, after which the luminescence response was quantified as described above.

### Calcium Mobilization Assay

Peptide conjugates were prepared as described for the tango assay. MDA-MB-231 cells were harvested and washed with PBS (pH 7.4). The MDA-MB-231 cells were then treated with 10 µM of either conjugate for 10 minutes at 37 °C at pH 7.4 or 6.0. After treatment, the MDA-MB-231 cells were washed once with PBS at the same pH as the treatment and resuspended in HBSS supplemented with 20 mM HEPES. NK-92 cells were loaded with Fluo-4 NW (Invitrogen) according to the manufacturer’s protocols. The cells were incubated for 30 minutes at 37 °C and then for 30 minutes at room temperature. The fluorescence emission at 516 nm (excitation at 494 nm) of 160,000 NK-92 cells was acquired with a Fluorolog-3 Spectrofluorometer (HORIBA) for approximately 30 seconds after which 40,000 MDA-MB-231 cells were added, and the fluorescence was monitored continuously.

## Acknowledgement

This work was supported by internal funds from Lehigh University to D.T., and National Institute of General Medical Sciences Grant R35GM124893-01 to M.M.P.

## Abbreviations

CD: Circular Dichroism.
FPR: Formyl peptide receptor.
LUV: Large unilamellar vesicles.
MALDI-TOF: Matrix-assisted laser desorption/ionization time of flight.
NK: Natural killer
pHLIP: pH(Low) Insertion Peptide.
POPC: 1-palmitoyl-2-oleoyl-sn-glycero-3-phosphocholine.
RP-HPLC: reverse-phase high-performance liquid chromatography.

## Notes

### Competing Interest Statement

The authors have declared no competing interest.

