## Supplementary Information for "Selective Display of a Chemoattractant Agonist on Cancer Cells Activates the Formyl Peptide Receptor 1 on Immune Cells"

**Scheme S1.** Synthesis of FPRL-pHLIP conjugates.

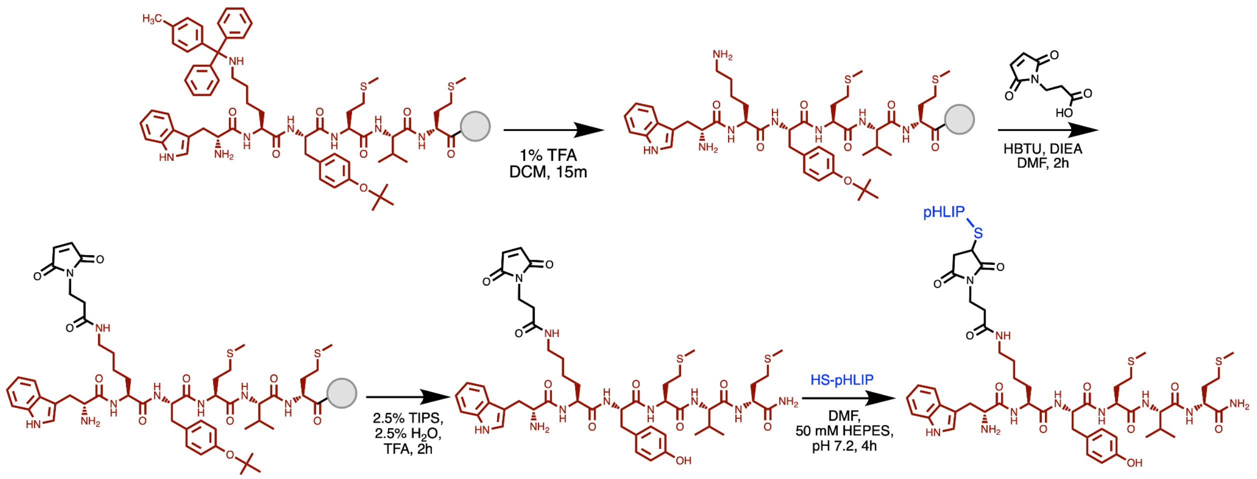

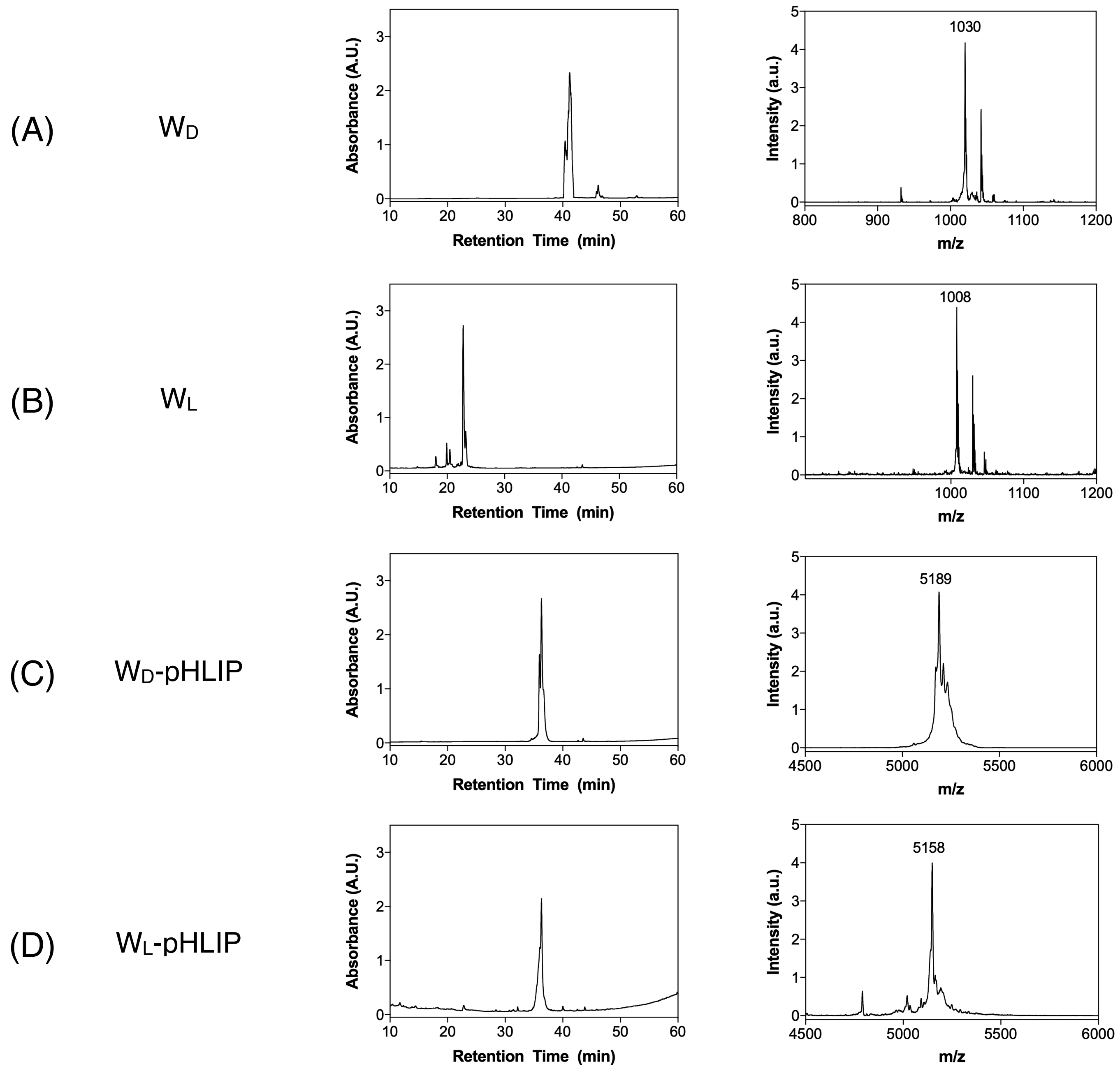


**Figure S1.** Purity check by RP-HPLC and MALDI-TOF MS spectra of synthesized peptides. **(A)** $W_D$: calculated ($MNa^+$) = 1031, found ($MNa^+$) = 1030. **(B)** $W_L$: calculated ($MH^+$) = 1008, found ($MH^+$) = 1008. **(C)** $W_D$-pHLIP: calculated ($MNa^+$) = 5182, found ($MNa^+$) = 5189. **(D)** $W_L$-pHLIP: calculated ($MH^+$) = 5159, found ($MH^+$) = 5158.

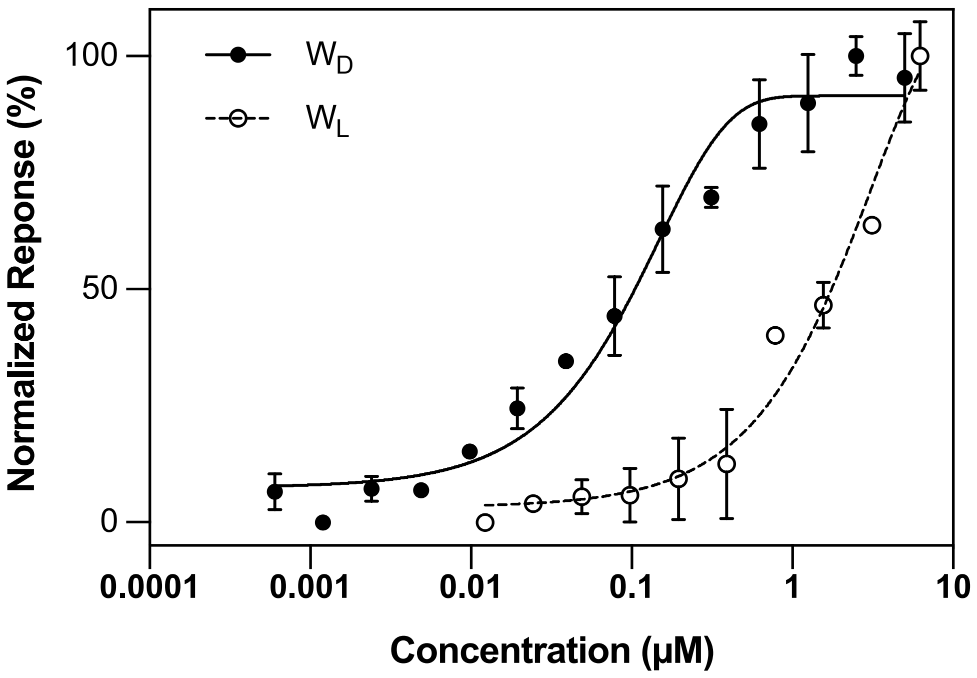


**Figure S2.** Concentration-dependent response of FPR1 activation by the $W_L$ (dashed line) or $W_D$ (solid line) peptides. Results are shown as mean ± standard error ($n$ = 3).
